# The Role of Morphological Adaptability in *Vibrio cholerae*’s Motility and Pathogenicity

**DOI:** 10.1101/2024.03.27.586043

**Authors:** Jun Xu, Keigo Abe, Toshio Kodama, Marzia Sultana, Denise Chac, Susan M. Markiewicz, Erika Kuba, Shiyu Tsunoda, Munirul Alam, Ana A. Weil, Shuichi Nakamura, Tetsu Yamashiro

## Abstract

*Vibrio cholerae*, the etiological agent of cholera, exhibits remarkable adaptability to different environmental conditions by undergoing morphological changes that significantly contribute to its pathogenicity and impact the epidemiology of the disease globally. This study investigates the morphological adaptability of the clinically isolated *V. cholerae* O1 strain, specifically focusing on the motility and pathogenicity differences between the filamentous and original comma-shaped forms within diverse viscosity conditions. Utilizing the El Tor strain of *V. cholerae* O1, we induced the transformation into the filamentous form and performed a comparative analysis with the canonical comma-shaped morphology. Our approach involved assessing motility patterns, swimming speeds, rotation rates, kinematics, and reversal frequencies through dark-field microscopy and high-speed imaging techniques. The findings reveal that filamentous *V. cholerae* cell retains enhanced motility in viscous environments. This suggests an evolutionary adaptation enabling survival across a range of habitats, notably the human gastrointestinal tract. Filamentous forms demonstrated increased reversal behavior at mucin interfaces, hinting at an advantage in penetrating the mucus layer. Rabbit intestinal loop assays further showed that both morphological forms exhibit similar fluid accumulation ratios, thus indicating comparable pathogenic potentials. These results underscore the significance of *V. cholerae*’s morphological flexibility in adapting to environmental viscosity changes, shedding light on the bacterium’s intricate survival and infection strategies. Our study provides critical insights into the dynamics of cholera, underlining the importance of considering bacterial morphology in developing effective cholera control strategies.

## INTRODUTION

*Vibrio cholerae*, a Gram-negative, curved rod, comma-shaped bacterium, the causative agent of cholera, is a significant global public health concern, particularly in the regions with limited access to clean water and sanitation (*1*). *V. cholerae* is responsible for severe diarrheal outbreaks, leading to high morbidity and mortality rates worldwide. The ongoing challenge of cholera, characterized by millions of cases annually, predominantly in under-resourced areas, underscores the critical need for a comprehensive understanding of this pathogen’s biology and mechanisms of disease (*2*). The impact of cholera extends beyond immediate health implications, affecting socio-economic stability and development. Rapid spread in communities, potential for severe dehydration, and high fatality rates in untreated cases make cholera a key target for public health interventions (*3*). The disease often highlights broader issues related to water safety and public health infrastructure, serving as an indicator of environmental and societal health (*4*).

At the molecular level, the virulence of *V. cholerae* is primarily attributed to the cholera toxin, which disrupts intestinal electrolyte balance, causing the characteristic watery diarrheal disease (*5*). Additionally, the bacterial motility, facilitated by a single polar flagellum, plays a critical role in its ability to colonize the human intestine and navigate through aquatic environments (*6*). This motility is multifaceted, contributing not just to locomotion but also to biofilm formation, environmental sensing, and host tissue interactions (*7–10*). The motility of *V. cholerae* is a complex and finely regulated process, involving a series of cellular and molecular mechanisms that enable the bacterium to respond to environmental cues and adapt to varying conditions.

Despite extensive research, *V. cholerae* continues to reveal complexities in its adaptation strategies. Under specific environmental conditions, it can undergo a dramatic morphological transformation into an elongated, helical filamentous form (*11*). This change raises important questions about its impact on bacterial motility and pathogenicity. The filamentous form of *V. cholerae* represents a significant deviation from its typical morphology, potentially altering its swimming behavior, colonization efficiency, and interactions with host tissues (*12*).

Our study is motivated by the need to understand this morphological adaptation and its consequences for *V. cholerae* motility and pathogenicity. Focusing on bacterial morphological versatility, this research aims to shed light on the implications of these changes for *V. cholerae* infection and persistence. We hypothesize that the filamentous form of *V. cholerae*, induced by environmental factors, may exhibit distinct motility patterns that are crucial in understanding the dynamics of cholera outbreaks. Through this investigation, we seek to contribute to the broader understanding of *V. cholerae*’s adaptive mechanisms. This research has implications for scientific understanding of bacterial motility and adaptability, and it holds potential for informing public health strategies to prevent and control cholera. By elucidating the nuances of *V. cholerae*’s morphological and motility changes, our goal is to provide insights critical for addressing this persistent global health challenge.

## RESULTS

### Induction of Filamentation of *V. cholerae* Cells

*V. cholerae* can change its cell morphology from comma-shaped to filamentous in response to various natural cues. To study the behavior and function of the filamentous form, we have successfully induced comma-shaped *V. cholerae* cells into their filamentous form using nalidixic acid. Importantly, contrary to what might be inferred, this morphological change did not result in the loss of motility. Instead, the filamentous cells retained the capability for movement, although the nature of their motility, specifically the speed and mechanisms of movement, underwent modifications reflective of their altered morphology. Our modified method, adapted from previous methodologies (*13*, *14*), resulted in over 80% of the cells transforming into the filamentous form within 6 to 18 hours (Fig1, Video S1). These cells retained their motility for several days, indicating effective induction without compromising cellular function.

**Figure 1:**
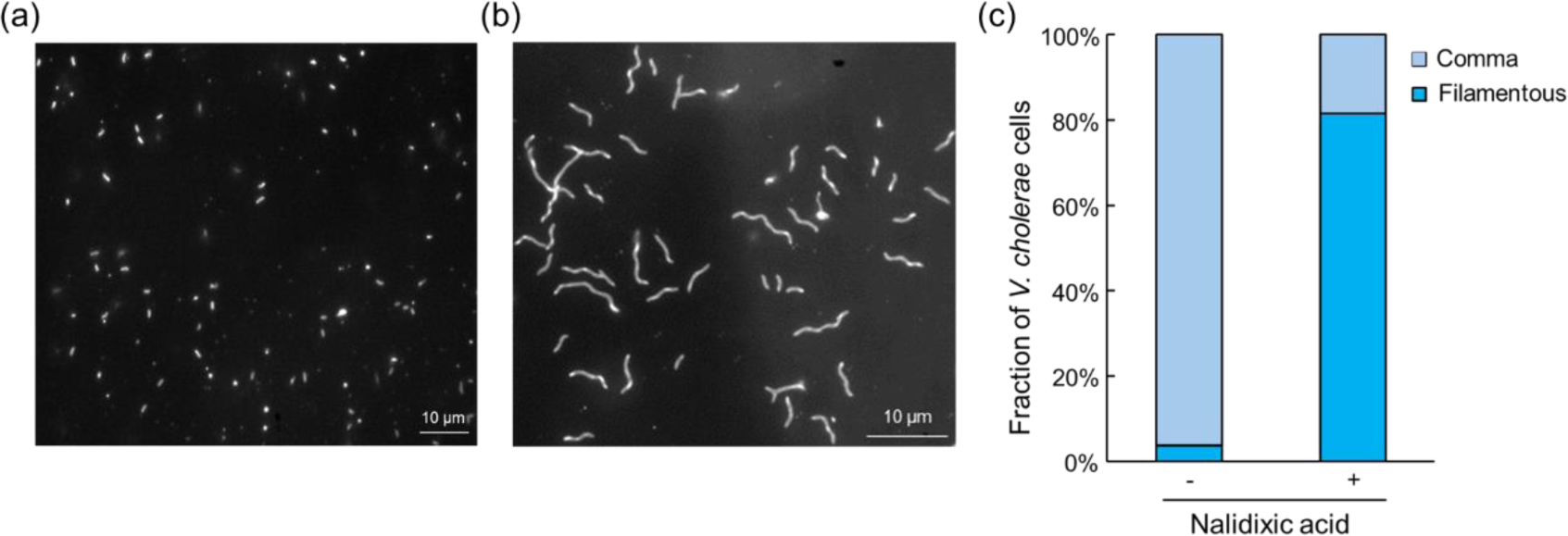
Induction of Filamentous *V. cholerae* Cells. Dark-field microscopic images illustrate the transformation of *V. cholerae* cells into their filamentous form. **(a)** displays the typical comma-shaped *V. cholerae* cells prior to induction. **(b)** shows the cells after undergoing 18 hours of induction with nalidixic acid. **(c)** presents the proportion of *V. cholerae* cells that transformed into filamentous forms in the induction solution, comparing conditions with and without nalidixic acid over an 18-hour period. The data is aggregated from a sample size of n=150 cells for each condition, compiled from three independent experimental trials.

### Swimming Patten of Filamentous *V. cholerae* Cells

Filamentous cells showed distinct swimming patterns compared to the typical ‘shooting star’ movement seen in comma-shaped cells (Video S2). While they lost the ability to perform the ‘flick’ movement typical of the latter, they adapted to turning maneuvers with large radius or “back and forth” movements, particularly when changing direction (Fig 2a, b). Fig. 2c demonstrates how the swimming patterns of *V. cholerae* cells are influenced by changes in the viscosity of their environment. In high viscosity environments, comma-shaped cells exhibited reduced travel distance (Fig. 2c left) and ‘flick’ frequency (Fig. S4), whereas filamentous cells’ movement patterns remained relatively unaffected (Fig 2c right).

**Figure 2:**
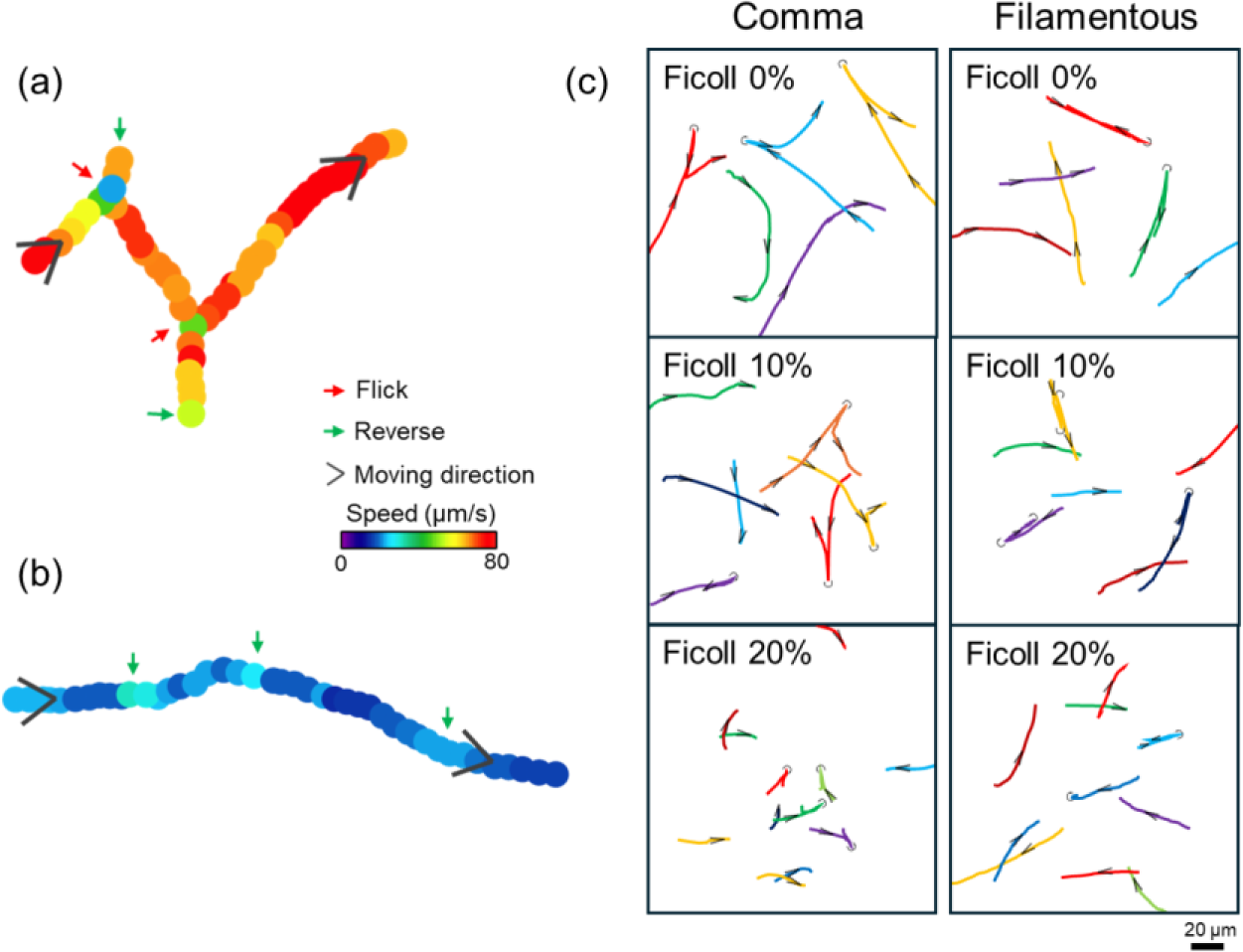
Motility pattern of *V. cholerae* Cells. **(a)** and **(b)** illustrate the swimming pattern of a typical comma-shaped cell and a typical filamentous cell, respectively. Red arrows highlight instances of ‘flicking’ movements—a rapid change in direction facilitated by the bacterium’s flagellum, with green arrows indicating moments of reversal in the cell’s trajectory. Black arrows represent the general direction of movement, while the variation in swimming speeds is conveyed through color coding, providing an intuitive visualization of speed differences. **(c)** displays the swimming trajectories over a 5-second period for representative *V. cholerae* cells in environments of varying viscosities. Black arrows and U-turn symbols illustrate the direction of movement and instances of reversal maneuvers, respectively.

### Swimming Kinematics of Filamentous *V. cholerae* Cells

In the medium with low viscosity such as water, filamentous cells swim at an average speed of 20 μm/s, slower than the 100 μm/s average in comma-shaped cells(Fig 3a), with a correlation measured between increased body length and decreased speed (Fig S5 black dots/line). Both cell types displayed viscosity-dependent motility, with dramatic impairments in comma-shaped cells at viscosities above 10 mPa•s (Fig 3a). Filamentous cells exerted significantly higher forces during locomotion compared to comma-shaped cells, as indicated by the propulsive force of approximately 0.2 pN for comma-shaped cells and nearly ten times that for filamentous cells (Fig 3b). This difference in force generation highlights the powerful locomotion of the filamentous form, highlighting the adaptability of the cells to these conditions. Furthermore, filamentous cells generated about 2000 pN nm of torque at a rotation rate of 10 Hz in motility buffer, with a linear decrease in torque as Ficoll concentration increased (Fig S6).

**Figure 3:**
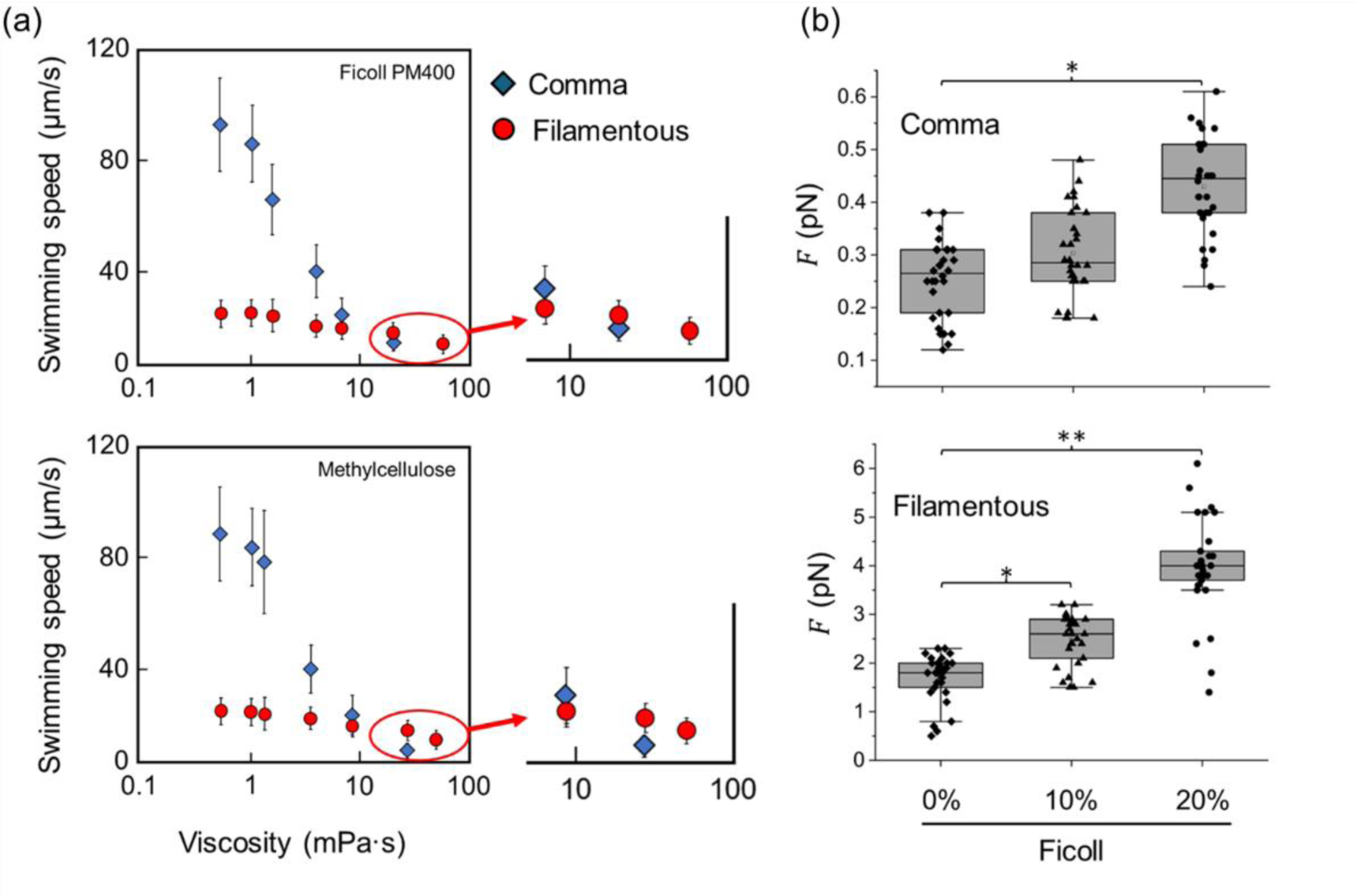
Kinematic Parameters of *V. cholerae* Cells. **(a)** presents the average swimming speeds of both comma-shaped and filamentous *V. cholerae* cells in media with varying viscosities. The enlarged insets provide a closer look at the bacterial motility in highly viscous environments. **(b)** The box charts that detail the force exerted on the two morphological forms of *V. cholerae* while swimming in media with different viscosities. A one-way ANOVA with Dunnett’s test was conducted to assess the statistical significance of the observed differences in force exertion, with asterisks indicating significant differences (\**P* < 0.05, \*\**P* < 0.01). The data is aggregated from a sample size of n=60 cells for each viscosity level, compiled from three independent experimental trials.

### Reversal Behavior During Swimming

To observe more detailed swimming behavior of these *V. cholerae* cells, we utilized a flow chamber setup, to mimic the natural aquatic environments of *V. cholerae*. The flow chamber consists of two main phases: a liquid phase and a mucin phase, which together create an interface that simulates the gradient encountered by *V. cholerae* at the boundary between aqueous environments and the more viscous mucus layers of the human intestine (Fig 4a). Filamentous *V. cholerae* cells displayed a unique reversal behavior at the mucin-liquid interface, characterized by distinct “back and forth” movements (Fig 4b and c, Video S3-2). This behavior sharply contrasts with that of the more agile comma-shaped cells, which often became entrapped at the same interface (Fig 4b and d, Video S3-1). The frequent and pronounced reversal movements of filamentous cells suggest an adaptive mechanism for efficient navigation in viscous environments. The “back-and-forth” motion demonstrated by these cells is depicted as a strategy to explore and potentially penetrate the viscous mucus barrier, seeking more accessible routes through this physical obstacle. This strategy potentially facilitates penetration of mucus layers, an essential component of the bacterium’s pathogenic process, as illustrated in Figure 4e.

**Figure 4:**
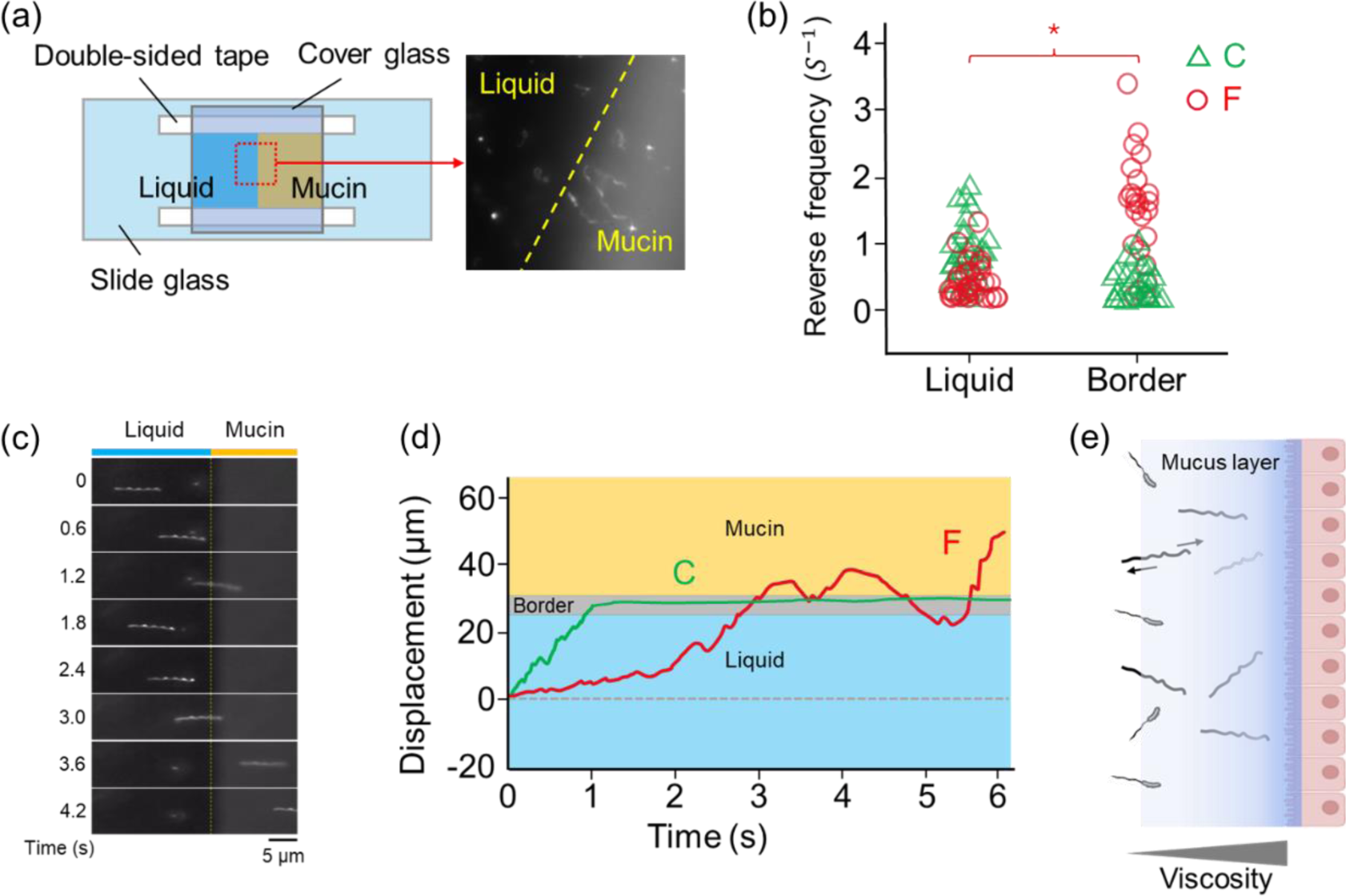
Reversal Behavior of Filamentous *V. cholerae* Cells. **(a)** A schematic representation of the flow chamber setup containing adjoining liquid and mucin phases. The initial positioning of bacterial cells in the liquid phase at the start of the experiment is shown. **(b)** A graph depicting the reversal frequency of comma-shaped (green triangle) and filamentous (red circle) cells in both the liquid phase and at the border. A Mann-Whitney U test revealed a significant difference in the reversal frequencies of filamentous cells between the liquid phase and the border area (\**P* < 0.05), with n=20 cells evaluated for each condition. **(c)** Time course of the locomotion of a filamentous cell observed at the liquid-mucin border. **(d)** Tracks illustrating the displacement of individual comma-shaped (indicated by ‘C’) and filamentous (indicated by ‘F’) cells as they swim in the liquid and at the border between the liquid and mucin phases. The grey band represents the border’s position. **(e)** A schematic illustrating how frequent reversal movements in filamentous cells contribute to their navigation.

### Comparative Infection Abilities of Filamentous and Comma-Shaped *V. cholerae*

In the rabbit intestinal loop assays, we found that infection with filamentous and comma-shaped *V. cholerae* cells displayed similar pathogenic capabilities. The fluid accumulation ratio (FAR), indicative of cholera-like symptoms, was almost identical for both morphologies, with filamentous cells showing an FAR of 1.1 mL/cm, closely aligning with that of the comma-shaped cells. This similarity in FAR suggests that the transformation into filamentous form does not impede the bacterium’s ability to induce comparable lesions in the host’s intestine (Fig 5a, b). This is further consistent with the unchanged expression of *ctxA* gene between the LB co-culture of comma-shaped and filamentous cells, confirmed by a qPCR analysis (Fig S7). Throughout the 18-hour infection period, a significant proportion of the cells maintained their injected morphological form, whether comma-shaped or filamentous (Fig 5c, d), underscoring the stability of filamentous shape within the host gut environment. It was also noted that filamentous cells were not discovered from intestinal loops that had been injected with comma-shaped *V. cholerae* (Fig 5e), suggests that, under the conditions of our experiment, the transformation from comma-shaped to filamentous morphology did not occur within the rabbit intestinal loop environment or was not detectable within the duration.

**Figure 5:**
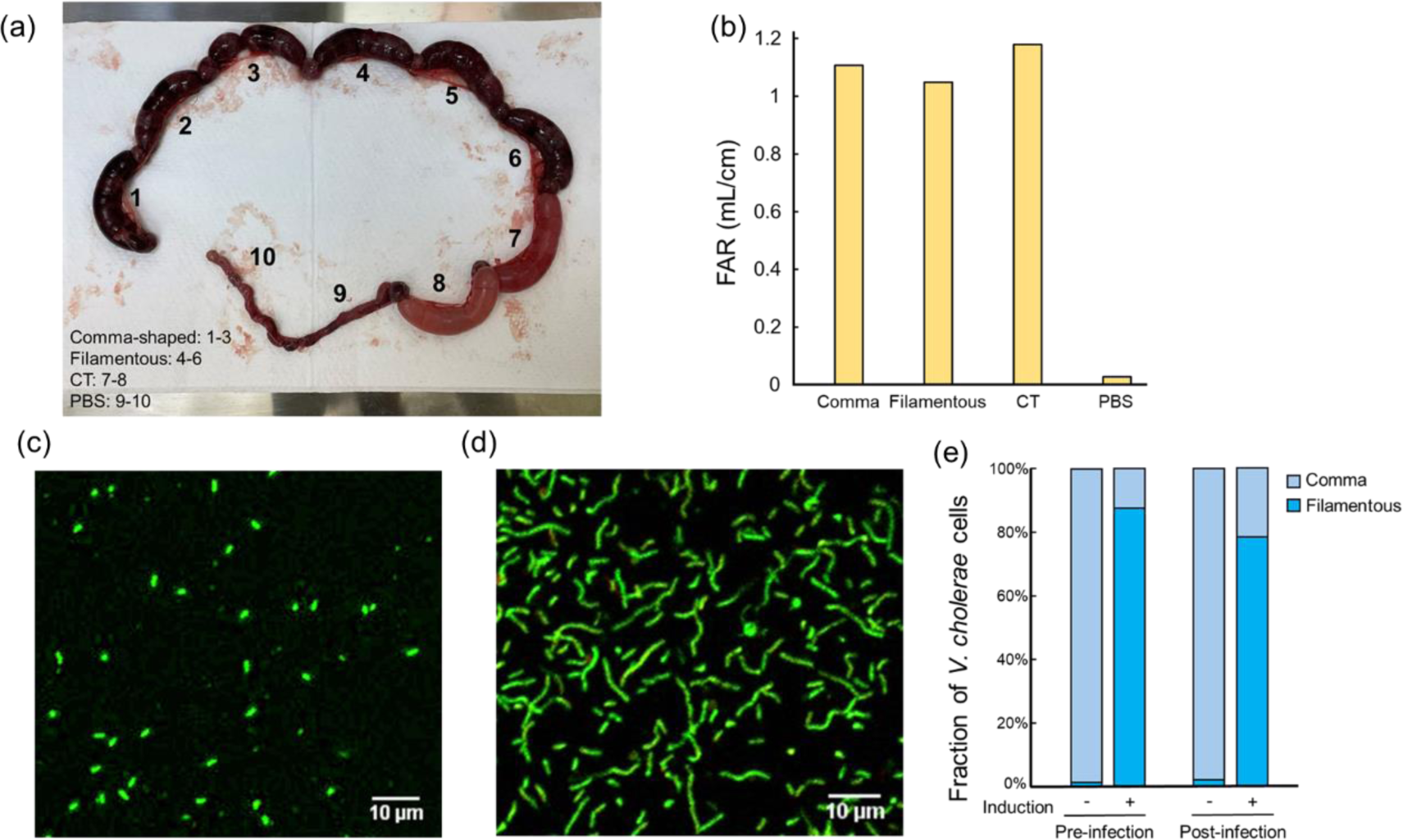
Comparative Infection Abilities of Two Morphologies of *V. cholerae*. **(a)** The image shows the intestinal loops 18 hours post-injection, clearly differentiating between loops injected with different agents. Loops 1-3 contain comma-shaped cells, loops 4-6 filamentous cells, loops 7-8 cholera toxin as a positive control, and loops 9-10 phosphate-buffered saline (PBS) as a negative control. The dark-black coloration in loops 1-6 is indicative of severe lesions induced by the injected *V. cholerae* cells. **(b)** Graph illustrating the fluid accumulation ratio (FAR) in the intestinal loops. This provides a comparative measure of pathogenicity between loops injected with *V. cholerae* cells in two morphologies, cholera toxin (CT), and PBS. The FAR values serve as an indicator of the severity of infection caused by each agent. **(c) and (d)** Epifluorescent microscopy images display intestinal fluid samples collected from rabbit intestinal loops 18 hours post-injection with comma-shaped **(c)** and filamentous **(d)** *V. cholerae* cells, respectively. **(e)** Proportion of *V. cholerae* cells pre- and post-infection period. The inoculum morphologies were found to be maintained after the 18 hours infection.

### Presence of Filamentous *V. cholerae* Cells upon exposure to Bile

When cultured in L-broth supplemented with bile, *V. cholerae* exhibited filamentation, though to a lesser extent than when induced by antibiotics (Fig 6). This observation hints at the possible natural formation of filamentous *V. cholerae* within the human small intestine. Further supporting this notion, direct fluorescent antibody (DFA) assays conducted on rice watery stool samples from patients with cholera showed the presence of filamentous *V. cholerae* O1 cells (Fig S8). These findings suggest a potential role for filamentous *V. cholerae* O1 in the pathogenesis of cholera, indicating that this morphological form could have significant implications in the natural course of the disease diarrhea.

**Figure 6:**
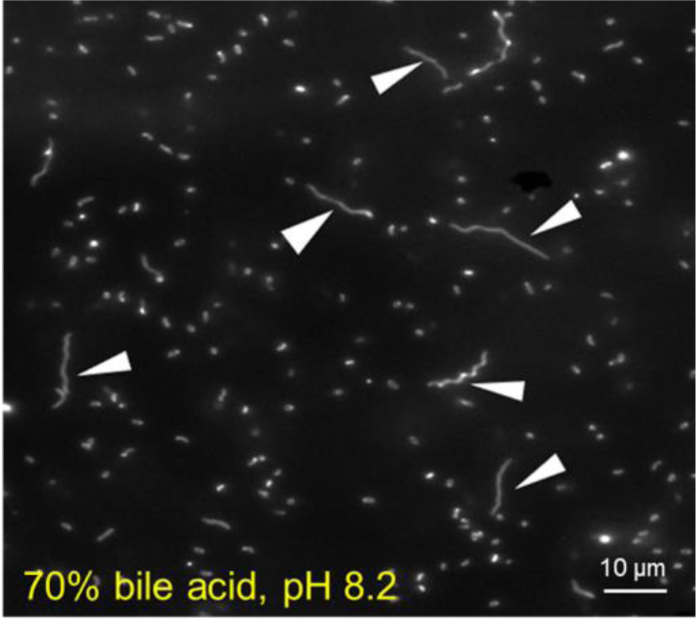
Presence of Filamentous *V. cholerae* Cells upon Exposure to Bile. Dark-field microscopic image that displays the filamentation of *V. cholerae* cells cultured in Luria-Bertani (LB) medium supplemented with bile. White arrows indicate the filamentous cells.

## DISCUSSION

*V. cholerae* remains a formidable public health challenge, particularly in regions with inadequate water and sanitation infrastructure. The adaptability and resilience of *V. cholerae*, notably in its ability to assume different morphological forms, have long been subjects of interest in microbiological research (*15*, *16*). Our study contributes significantly to this field by elucidating the differences in motility and pathogenicity between filamentous and comma-shaped forms of *V. cholerae*, particularly under varying environmental conditions. Our findings reveal that the filamentous form of *V. cholerae*, despite its reduced swimming speed, exhibits enhanced motility in viscous environments. This suggests an evolutionary adaptation that enables the bacterium to navigate and survive in diverse habitats, including the human gastrointestinal tract, a key site for *V. cholerae* infection (*17*, *18*). The ability of filamentous cells to perform frequent reversal movements at mucin borders is a novel insight, potentially facilitating mucus layer penetration, an essential factor in the pathogenesis of cholera (*19*, *20*). Together these data suggest an enhanced ability of the filamentous *V. cholerae* for penetration and travel through the mucous layer, which may confer an advantage over the solitary comma-shaped cell to cross the thick mucin layer for reaching the small intestinal gut epithelium (Fig 4e).

In terms of pathogenicity, our study underscores that morphological transformation does not compromise *V. cholerae*’s disease-causing ability, as evidenced by the comparable fluid accumulation ratios between filamentous and comma-shaped forms. This observation suggests that filamentous cells retain their potential to induce cholera-like symptoms, consistent with studies that have emphasized the role of bacterial morphology in infection dynamics (*10*, *21*). However, this raises an intriguing point of discussion regarding the specific advantages of filamentous motility and their manifestation in pathogenicity. Given the enhanced ability of filamentous cells to navigate through viscous zones such as mucus, we might intuitively expect an increase in pathogenicity for these morphological forms. Yet, the direct injection of bacterial fluid into the rabbit intestinal loops may not fully capture the nuances of a natural infection process that typically occurs through ingestion of contaminated food and water. The advantages of filamentous motility may not be discernible using the rabbit loop model, which bypasses the initial stages of infection encountered in food-borne transmission and entry through the mouth. Therefore, although no difference in the disease-causing ability was observed in this experiment, the virulence of filamentous cells compared to comma-shaped cells requires further investigation, explore if there are differences in their ability to reach the gut epithelial surface or variations in the expression of virulence factors to cause disease.

Crucially, the presence of filamentous *V. cholerae* in cultural medium with bile and clinical stool samples from patients with cholera confirms the relevance of these morphological forms in natural infection scenarios, echoing findings from previous clinical studies (*22*, *23*). This correlation between laboratory and clinical observations not only underscores the filamentous form’s role in the natural course of *V. cholerae* infections but also suggests its potential contribution to the spread of the disease. The filamentous morphology may offer several advantages that facilitate the *V. cholerae* infection. Firstly, the enhanced ability of filamentous cells to navigate through viscous environments, such as mucus layers, could allow for more efficient colonization of the host’s intestinal tract, increasing the likelihood of successful infection and subsequent shedding into the environment. Additionally, the physical robustness and increased surface area of filamentous cells may potentially enhance adherence and better survival on various surfaces, including food and water sources, thereby increasing the opportunities for ingestion and infection of new hosts. Furthermore, the filamentous form may possess altered immunogenic properties or evade host immune responses more effectively than the comma-shaped form, contributing to sustained infection rates and enhancing transmission dynamics. Lastly, the presence of filamentous *V. cholerae* O1 cells in rice watery stool of patients hospitalized at icddr,b Dhaka hospital suggests that these forms could shed into the environment where they could circulate in the aquatic environment, potentially transforming back to the comma-shaped form under suitable conditions for maintaining cholera transmission cycle during epidemic periods. Thus, these cells could persist in the clusters of biofilms in their aquatic reservoirs when the epidemic is over. This cycle of morphological transformation and environmental persistence underscores a complex strategy for survival and transmission, highlighting the need for further research to elucidate the specific mechanisms through which filamentous *V. cholerae* contributes to the environmental adaptation of *V. cholerae* and the spread of cholera (*24*, *25*).

It is also imperative to consider the morphological stability of *V. cholerae* throughout the course of an infection. The fact that filamentous cells were not collected from the rabbit ileal loops injected with comma-shaped *V. cholerae*, alongside the identification of filamentous forms in human cholera stools, warrants further discussion about the *V. cholerae*’s ability to maintain its form during the course of its infection. This observation is crucial, as it hints at a potential morphological consistency that *V. cholerae* exhibits from ingestion to excretion. Notably, the presence of filamentous *V. cholerae* in human stool samples suggests the potential role of this form in cholera pathogenesis; although, it remains unclear whether patients initially ingested the bacterium in its filamentous form, or it was induced from comma-shaped morphology during the course of the infection. Although there is a remote possibility that *V. cholerae* preserves its cellular morphology throughout the infection process in humans, our experiment elucidates those environmental conditions, notably exposure to antibiotics and bile, that induce the morphological transition of *V. cholerae*. This complements the observations that *V. cholerae* can modulate its cellular morphology, adopting a coccoid form as a strategic response to mitigate stress or unfavorable environmental conditions (*26*). Thus, the emergence of filamentous morphology could be interpreted as another strategic adaptation by *V. cholerae* to navigate environmental stressors, including bile salts, simultaneously maintaining its pathogenic potential. Nevertheless, our experimental setup may not have fully mimicked the intricate conditions of the human intestine that trigger such morphological changes. This underscores the complexity of *V. cholerae*’s life cycle and the need for further investigation into how environmental factors and the bacterium’s inherent biological responses dictate its morphology during infection. The exploration of whether *V. cholerae* undergoes transformation or maintains its form across different stages of infection is paramount for understanding its pathogenic mechanisms and implications for disease spread (*27*, *28*).

In conclusion, our research provides pivotal insights into *V. cholerae*’s morphological flexibility and its implications for bacterial motility and pathogenicity. These findings enhance our comprehension of this pathogen and could inform future strategies for managing and controlling cholera outbreaks. Moving forward, further research should aim to unravel the molecular mechanisms underlying *V. cholerae*’s morphological transformations and explore their impact on interactions with the host immune system. Understanding these aspects in greater depth will be crucial for developing more effective interventions against this enduring public health challenge.

## MATERIALS AND METHODS

### *V. cholerae* Strain, Culture Medium, and Reagents

The *V. cholerae* O1 El Tor strain N16961 (ATCC 39315), known for its well-characterized flagellar and chemotactic systems, was selected for this study to accurately assess changes in motility. Bacterial cultures were grown in Luria-Bertani (LB) broth at 37 °C, maintaining a pH of 7.5. The LB medium comprised 10 g of tryptone, 5 g of yeast extract, and 5 g of NaCl per liter (Nacalai Tesque, Kyoto, Japan). For observing motility, *V. cholerae* cells were resuspended in TMN (Tris-MgCl2-NaCl) buffer, with a composition of 50 mM Tris-HCl, 5 mM MgCl2, 5 mM glucose, 100 mM KCl, and 200 mM NaCl at pH 7.5 (*29*). To simulate environmental conditions affecting motility, various reagents such as methylcellulose, Ficoll, bile acid, and mucin were introduced (all from Wako Pure Chemical, Osaka, Japan).

### Induction of *V. cholerae* into Filamentous Form

For filamentation, we used a solution containing 25 mg/mL yeast extract and 4 mg/mL nalidixic acid in artificial sea water (Sigma-Aldrich, MO, USA) (*13*). This solution was stored at 4°C to preserve its effectiveness. A single colony of *V. cholerae* was inoculated into 5 mL of fresh LB broth and incubated at 37°C with shaking until it reached an optical density at 600 nm (OD_600_) of 0.5-0.6. The cells were then centrifuged, resuspended in the induction solution, and incubated at 20°C in the dark without shaking. Morphological changes to filamentous form were monitored hourly using dark-field microscopy, with filamentous cells displaying intact motility typically observed after 6-18 hours of incubation.

### Fluorescent Staining and Microscopy Observation

Comma-shaped and filamentous *V. cholerae* cells were stained using fluorescein isothiocyanate (FITC). Fixed cells in 4% paraformaldehyde were washed, stained with FITC for one hour at room temperature in darkness, then mounted on glass slides using SlowFade Diamond anti-fade mounting medium (Life Technologies, CA, USA). Slides were examined under an epifluorescent microscope (BZ-X810, Keyence, Osaka, Japan) with standardized settings for exposure time and illumination.

### Observation of Bacterial Motility

Bacterial motility was observed using a dark-field microscope (Nikon Eclipse Ci-L Plus, Nikon, Tokyo, Japan) with one-sided illumination (*30*). Cell movement was recorded at 120 fps using a high-speed CMOS camera (Imaging Source, Taipei, ROC). Videos were analyzed quantitatively using ImageJ (NIH, MD, USA) and Excel (Microsoft, WA, USA), following established methodologies (*31–33*).

### Kinematic parameters of swimming *V. cholerae* cell

The geometric parameters of the *V. cholerae* cell bodies were measured and analyzed using ImageJ software. Modified resistive force theory and Stoke’s law were applied to estimate the swimming force for the comma-shaped and filamentous cells and the torque generated by the rotation of the filamentous cell body (*34–37*). The estimated swimming force was calculated based on the drag force acting on the cell body. The force required for swimming, denoted as *F_swim_*, is approximated to be equal to the total drag force, *F_drag_*, for straight-line swimming:

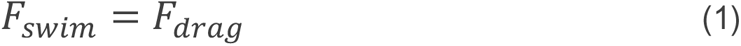

The total drag force *F_drag_* along the comma-shaped cell body:

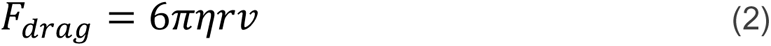

Equation (2) is the drag force on swimming comma-shaped cell, where *η* is the viscosity of the medium, *r* is the radius of the bacterium, and *v* is the swimming speed.

The total drag force *F_drag_* was derived by integrating different drag forces, *dF*, along the filamentous cell body:

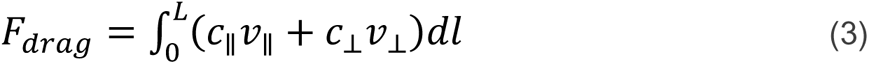

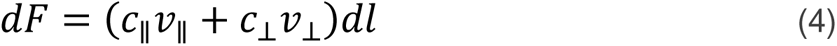

Equation (3) is the drag force on the swimming filamentous cell. In this equation, *c*_∥_ and *c*_⊥_ represent the parallel and perpendicular drag coefficient for motion, respectively. The velocity components, *v*_∥_ and *v*_⊥_, represent the translational and rotational speeds of each segment of the helical cell body, here we used the measured values of the swimming speed and rotation rate, respectively. The drag coefficients and the velocity components were calculated from the geometric parameters of the cell and the viscosity of the medium:

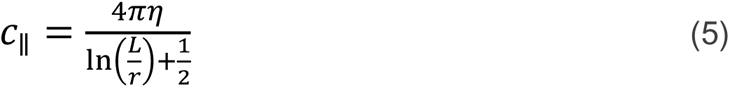

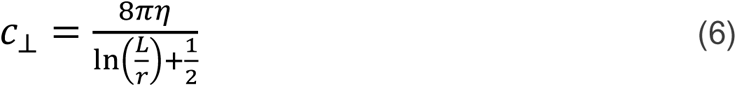

The length (*L*) and radius (*r*) of the cell body, were all determined (Table S2). The viscosity of the medium (*η*) was measured using a tuning-fork viscometer (SV-10, A&D company, MI, USA), as listed in Table S1.

The total torque exerted by the rotation of the cell was calculated by integrating the differential torque along the length of the filamentous cell body:

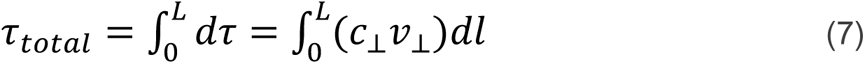

### Reversal Behavior During Swimming

To observe the reversal behavior of filamentous *V. cholerae* during swimming, a suspension of these cells was introduced into a flow chamber (Figure 4a) containing 5% mucin, creating a contiguous area of mucin and liquid with a distinct border between the two phases (*38–40*). This setup was previously developed as performed by Abe et al (2020), allowing for the detailed observation of bacterial movement at the interface of the two phases. The movement of the bacterial cells at this border area was closely monitored using a dark-field microscope (Nikon Eclipse Ci-L Plus, 40×/0.75, Nikon, Tokyo, Japan). The observed movements were recorded using a CMOS camera at a frame rate of 90 fps, for further detailed analysis.

### Rabbit intestinal loop assay

Comma-shaped and filamentous *V. cholerae* were cultured in Luria-Bertani broth at 37°C to mid-exponential phase, harvested, and resuspended in phosphate-buffered saline to 1 x 10^6^ cell/mL. New Zealand White rabbits (2 kg, 8 weeks old), fasted for 24 hours, underwent a surgical procedure under 2.5% thiopental (intravenous injection, 30 mg/kg) (NIPRO, Osaka, Japan) and 2.5% Isoflurane (Inhalation) (Wako Pure Chemical, Osaka, Japan) anesthesia to create intestinal loops. 2.5 mL of 2% Lidocaine (SANDOZ, Tokyo, Japan) was used for the local anesthetic to relieve pain. Segments of the small intestine were ligated into loops and inoculated with 1 mL of 10^6^ CFU/mL bacterial suspensions, cholera toxin, and sterile PBS for controls. After 18 hours, rabbits were euthanized with the intravenous injection of 10% thiopental, loops were harvested, and fluid accumulation was measured. Fluid accumulation ratios (FAR) were calculated and analyzed to assess the lesion induced by both forms of *V. cholerae* cells. Morphology of bacterial cells was further investigated by the fluorescent staining.

### *CtxA* Gene expression measurement

Briefly, induction of filamentous *V. cholerae* was performed using 15 ug/mL ampicillin (Fisher Bioreagents, PA, USA). Following incubation at the specified time points, samples underwent centrifugation and were resuspended in 1 ml TRIzol (Invitrogen, MA, USA) for RNA extraction, utilizing the Qiagen RNeasy Kit for extraction and the TURBO DNA-free Kit (Ambion, TX, USA) for DNA digestion, in accordance with the manufacturers’ instructions. cDNA synthesis was conducted using the High Capacity cDNA Reverse Transcription Kit (Applied Biosystems, MA, USA), and qPCR analysis was carried out on two 384-well plates employing PowerUp SYBR Green Master Mix (Applied Biosystems) on QuantStudio 6 Flex Real-Time PCR system (Applied Biosystems). Primers used for analysis are as follows: groEL forward 5’-ATGATGTTGCCCACGCTAGA-3’, groEL reverse 5’-GGTTATCGCTGCGGTAGAAG-3’ (*41*), ctxA forward 5’-TTGGAGCATTCCCACAACCC-3’, and ctxA reverse 5’-GCTCCAGCAGCAGATGGTTA-3’ (*42*). Data analysis was executed using PRISM GraphPad v10.1.2, with statistical significance assessed via 1-way ANOVA on relative expression levels (Livak Method). The housekeeping gene groEL served as a control in qPCR experiments.

### Statistical analysis

Statistical significance was analyzed using one-way analysis of variance (ANOVA) with Dunnett’s test, Mann-Whitney U test, as indicated in the figure legends (OriginPro 2023). \**P* < 0.05; \*\**P* < 0.01.

## SUPPLEMENTARY MATERIALS

Table S1. Viscosity of the medium used in the study.

Table S2. Geometric parameters of filamentous *V. cholerae* cells

Figure S3. Rotation rate of swimming filamentous *V. cholerae* cells

Figure: S4. Flicking frequency of comma-shaped *V. cholerae* cell.

Figure S5. Correlation between swimming speed, force, and cell length.

Figure S6. Torque-speed relation of swimming filamentous *V. cholerae* cells.

Figure S7. Gene expression of *ctxA* in two morphologies of *V. cholerae*.

Figure S8. Presence of filamentous *V. cholerae* cells in watery stool samples from patient with cholera.

Video S1. Swimming filamentous *Vibrio cholerae* cells.

Video S2. Example of swimming filamentous cells under one-sided illumination dark-field microscopy.

Video S3-1. Movement of the comma-shaped cells at the liquid-mucin border.

Video S3-2. Movement of the filamentous cell at the liquid-mucin border.

## Ethics Statement

The rabbit intestinal loop assay in this study adhered to the guidelines and received approval from the Institutional Animal Care and Use Committee at the University of the Ryukyus (Approval of research protocol: A2022050) and followed the principles of the Guide for the Care and Use of Laboratory Animals and Japan National Animal Welfare Laws. Efforts were made to minimize animal suffering, in alignment with the 3Rs principle. Rabbits were acclimatized, given ad libitum access to food and water, and housed to promote well-being before being anesthetized using thiopental, Lidocaine, and Isoflurane Inhalation to ensure minimal stress for the intestinal loop assay. Euthanasia was conducted humanely according to JAZA Guidelines on Euthanasia to ensure minimal suffering.

This study involves the analysis of watery stool samples from patients with cholera for the assessment of *V. cholerae* presence, was conducted under the ethical guidelines of the International Centre for Diarrhoeal Disease Research, Bangladesh (icddr,b). All procedures were approved by the icddr,b Ethical Review Committee (Approval of research protocol: # PR-23040). Informed consent was obtained from all participants or their legal guardians, ensuring understanding of the study’s purpose and the participants’ rights. Samples and data were anonymized and handled with strict confidentiality.

## Acknowledgment

We thank Naomi Higa (Univ. Ryukyus), Claudia Toma (Univ. Ryukyus), Hideyuki Matsunami (OIST), Matthias Wolf (OIST), and Tahmeed Ahmed (icddr,b) for the technical support and the helpful comments.

## Funding

This study was supported by the NIAID-AMED U.S.-Japan Cooperative Medical Sciences Program Collaborative Awards (22jk0210041h0001), the JSPS KAKENHI (20K22784), the Uruma Foundation for the Promotion of Science, and the Ryukyu Medical Association Award.

## Author contributions

Conceptualization of this study was primarily developed by JX, TK, MS, AAW, SN, MA, and TY. The formal experimentation, data collection and analysis were contributed by JX, KA, TK, MS, DC, SMM, AAW, EK, and ST. Software development and handling were conducted by SN, KA, and JX. Writing, review, and editing of the manuscript were jointly handled by JX, SN, AAW, MA and TY. Funding acquisition was contributed by JX, MS, AAW, MA and TY.

## Competing interests

The authors declare that they have no competing interests.

## Data and materials availability

All data needed to evaluate the conclusions in the paper are present in the paper and/or the Supplementary Materials. Additional data related to this paper may be requested from the authors.

## Notes

### Competing Interest Statement

The authors have declared no competing interest.

