## Supplemental materials for "The Role of Morphological Adaptability in *Vibrio cholerae*’s Motility and Pathogenicity"

### Supplementary Materials

Table S1. Viscosity of the medium used in the study.

Table S2. Geometric parameters of filamentous *V. cholerae* cells

Figure S3. Rotation rate of swimming filamentous *V. cholerae* cells

Figure: S4. Flicking frequency of comma-shaped *V. cholerae* cell.

Figure S5. Correlation between swimming speed, force, and cell length.

Figure S6. Torque-speed relation of swimming filamentous *V. cholerae* cells.

Figure S7. Gene expression of *ctxA* in two morphologies of *V. cholerae*.

Figure S8. Presence of filamentous *V. cholerae* cells in watery stool samples from patient with cholera.

Video S1. Swimming filamentous *Vibrio cholerae* cells.

Video S2. Example of swimming filamentous cells under one-sided illumination dark-field microscopy.

Video S3-1. Movement of comma-shaped cell at the liquid-mucin border.

Video S3-2. Movement of filamentous cell at the liquid-mucin border.

| Viscosity of medium | Symbol | Values |
|---|---|---|
| Without Ficoll or MC | $\eta$ | 0.87 mPa•s |
| With 5% Ficoll | | 1.88 mPa•s |
| 10% Ficoll | | 3.89 mPa•s |
| 15% Ficoll | | 7.58 mPa•s |
| 20% Ficoll | | 27.3 mPa•s |
| 25% Ficoll | | 48.0 mPa•s |
| 30% Ficoll | | 69.7 mPa•s |
| With 0.1% MC | | 1.23 mPa•s |
| 0.2% MC | | 2.32 mPa•s |
| 0.5% MC | | 6.44 mPa•s |
| 1% MC | | 32.9 mPa•s |
| 1.5% MC | | 84.7 mPa•s |

**Table S1. Viscosity of the Medium Used in the Study.** MC=methylcellulose

| Geometric parameters | Symbols | Values (2σ) |
|---|---|---|
| - Comma-shaped | | |
| Radius of cell body | $r$ | 0.25-0.4 μm |
| - Filamentous | | |
| Length of cell body | $L$ | 7.0-12.2 μm |
| Radius of cell body | $r$ | 0.31-0.4 μm |

**Table S2: Geometric Parameters of Filamentous *V. cholerae* Cells.** The geometric parameters were measured from a sample of 60 comma-shaped and 60 filamentous *V. cholerae* cells. Values falling within two standard deviations (2σ) of the mean are highlighted, representing the range that encompasses approximately 95% of all measured values, assuming a normal distribution. This approach provides a comprehensive overview of the typical geometric characteristics of *V. cholerae* cells under the study conditions.

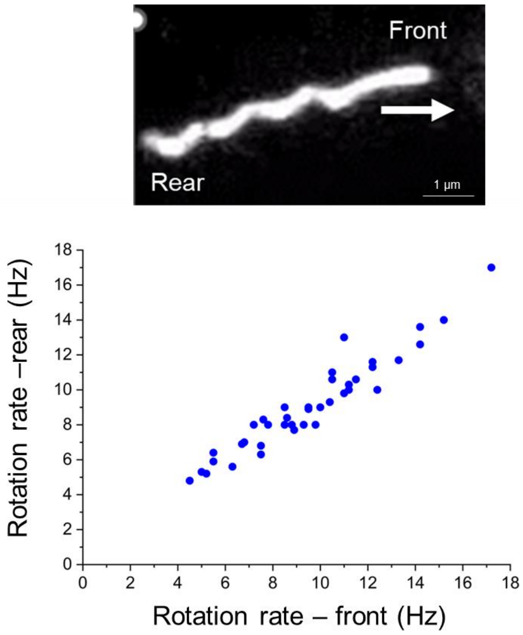


**Figure S3.** Rotation rates at the front and rear ends of the filamentous cells, with an accompanying image demonstrating these ends in a swimming filamentous cell; the arrow indicates the swimming direction.

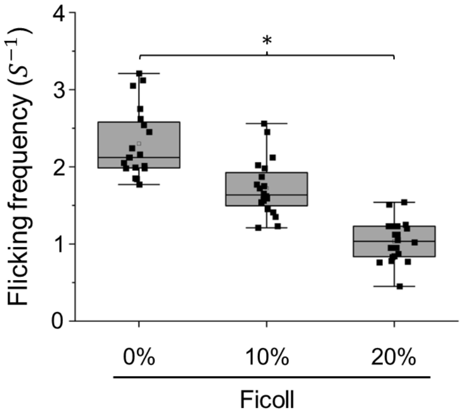


**Figure S4. Flicking frequency of comma-shaped *V. cholerae* cell.** The box chart details the change on flicking frequency in various viscosity, which decreases significantly in higher viscosity. A one-way ANOVA with Dunnett's test was conducted to assess the statistical significance of the observed difference, with asterisks indicating significant differences (*$P$ < 0.05, **$P$ < 0.01).

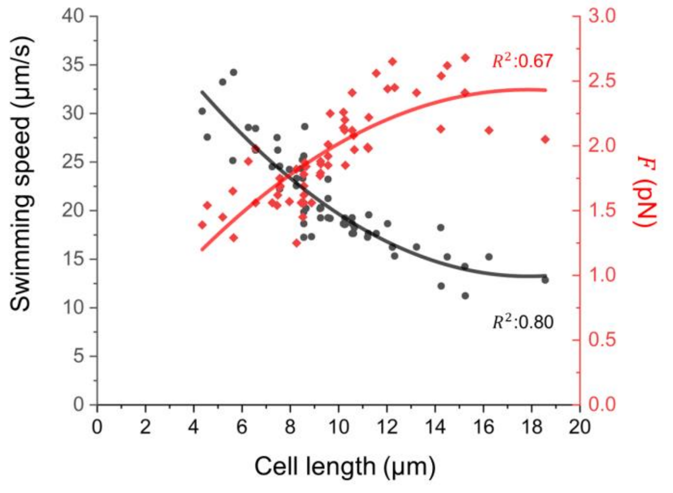


**Figure S5. Swimming Speed and Force of Swimming Filamentous *V. cholerae* Cells Correlated with Cell Length in low viscosity.** Graph displays the relationship between cell length and two parameters: swimming speed (black dots) and the force exerted (red dots) on filamentous cells swimming in a medium without Ficoll or methylcellulose. The black and red curves represent regression analyses for cell length-swimming speed and cell length-force (*F*), respectively. The data were collected from three independent experimental trials.

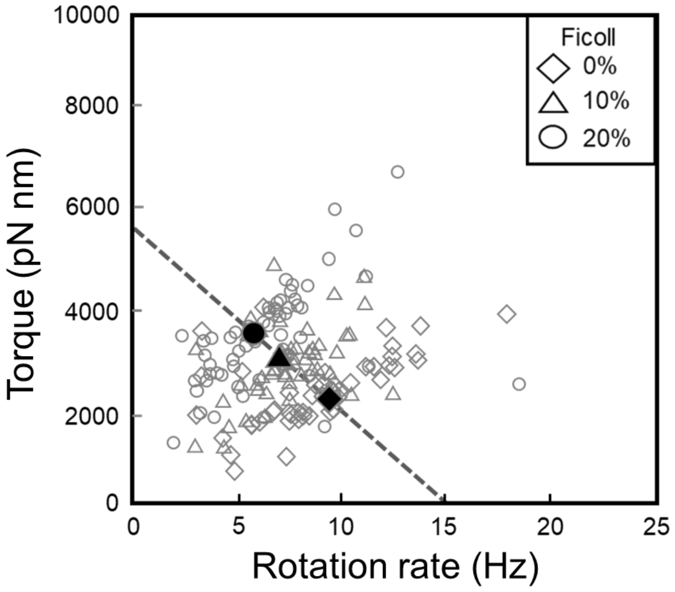


**Figure S6. Torque-Speed Relation of Swimming Filamentous *V. cholerae* Cells.** Graph shows the relationship between torque and rotation rate of the filamentous cell body in different viscosity conditions. Average values for each condition are presented in solid markers, with a fitted regression line (dashed line) illustrating the trend.

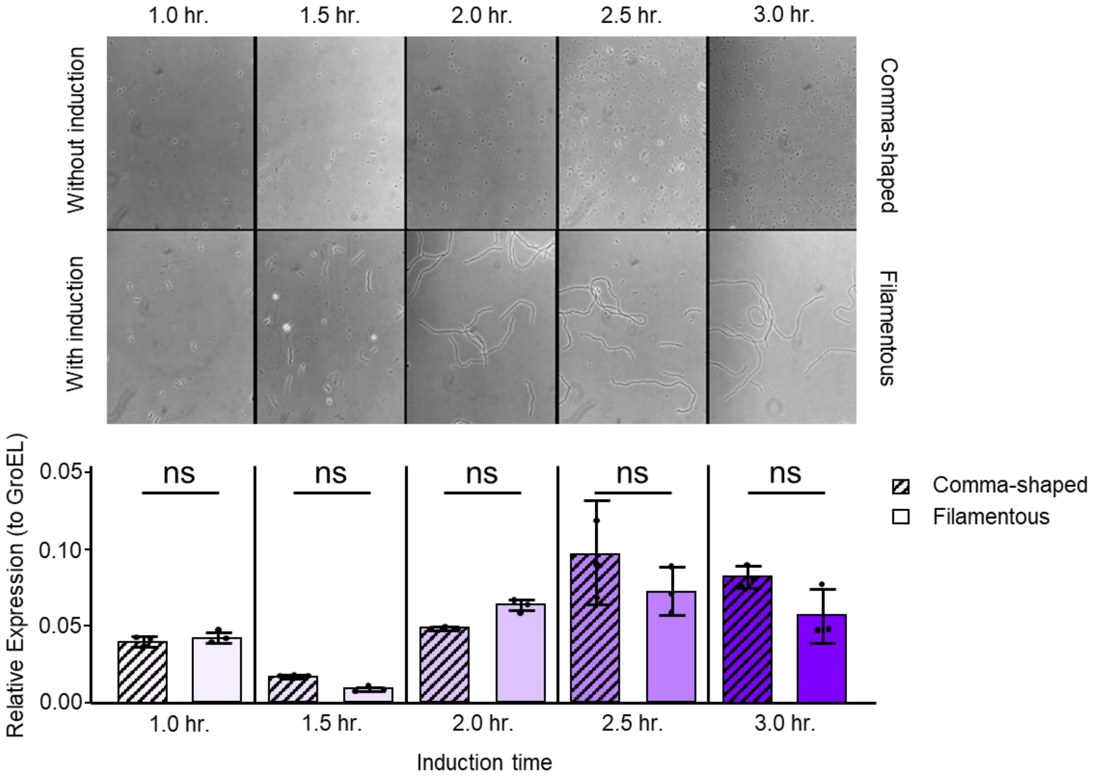


**Figure S7: Gene Expression of *ctxA* in Two Morphologies of *V. cholerae*.** This figure presents the time-course comparative analysis of *ctxA* gene expression between comma-shaped and filamentous *V. cholerae* in LB culture. Morphologies were confirmed using phase microscopy on a microscope (Zeiss AxioStar Plus, Carl Zeiss, Oberkochen, Germany) with 400X magnification at each time point post-induction. Images (upper) are representative of the phenotypes observed during the assay. The qPCR results (lower) indicate no difference (ns) in the transcriptional levels of the *ctxA* gene between the comma-shaped and filamentous forms of *V. cholerae*.

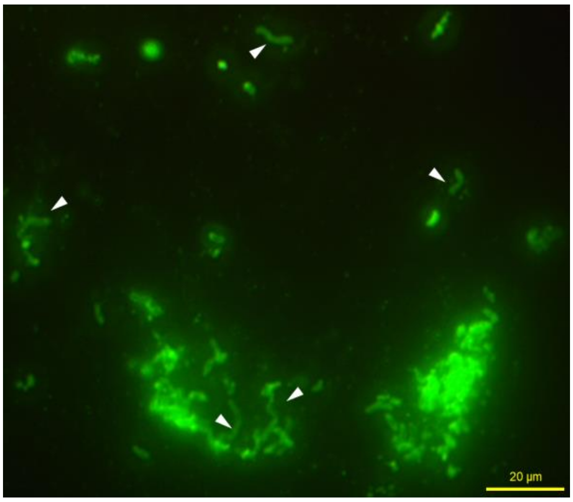


**Figure S8. Epifluorescent microscopy image of a direct fluorescent antibody (DFA) assay conducted on watery stool samples from patient with cholera hospitalized at icddr,b Dhaka hospital.** *V. cholerae* cells are labeled with a monoclonal antibody specific to *V. cholerae* O1, conjugated with fluorescein isothiocyanate (FITC), emitting a green fluorescence. White arrows point to the filamentous cells, indicating their presence in clinical samples of cholera stool.
